# The role of TORC1 in muscle development in Drosophila

**DOI:** 10.1101/010991

**Authors:** Isabelle Hatfield, Innocence Harvey, Erika. R. Yates, JeAnna R. Redd, Lawrence T. Reiter, Dave Bridges

**Author notes:** Corresponding Author: Department of Physiology, 894 Union Ave, University of Tennessee Health Science Center, Memphis, TN, 38138.

## Abstract

Myogenesis is an important process during both development and muscle repair. Previous studies suggest that mTORC1 plays a role in the formation of mature muscle from immature muscle precursor cells. Here we show that gene expression for several myogenic transcription factors including *Myf5*, *Myog* and *Mef2c* but not *MyoD* and myosin heavy chain isoforms decrease when C2C12 cells are treated with rapamycin, supporting a role for mTORC1 pathway during muscle development. To investigate the possibility that mTORC1 can regulate muscle *in vivo* we ablated the essential dTORC1 subunit *Raptor* in *Drosophila melanogaster* and found that muscle-specific knockdown of *Raptor* causes flies to be too weak to emerge from their pupal cases during eclosion. Using a series of GAL4 drivers we also show that muscle-specific *Raptor* knockdown also causes shortened lifespan, even when eclosure is unaffected. Together these results highlight an important role for TORC1 in muscle development, integrity and function in both Drosophila and mammalian cells.

## Background

The mTOR signaling pathway plays important roles during development in all eukaryotes and mTORC1 is a critical nutrient sensing protein kinase conserved in all eukaryotic organisms^1, 2^. This kinase responds to nutrient and growth hormone signals in the environment and subsequently phosphorylates targets involved in aging, growth, protein lipid and glycogen metabolism^3–5^. In addition to these effects on differentiated cells, there is an emerging role for mTORC1 in the regulation of cellular differentiation during development including neurogenesis^6, 7^, adipogenesis^8^ and myogenesis^9–11^. Consistent with these findings, either loss of the obligate mTORC1 complex members mTOR and Raptor, or treatment with rapamycin induces developmental lethality in mice^12–14^, worms^15^ and fruit flies^16^.

New muscle fiber formation occurs via the differentiation of muscle precursor cells called satellite cells^17, 18^. This process involves a cascade of transcription factors including several basic helix-loop-helix transcription factors such as *Myf5, Myog, Myod* and *Mef2c* (reviewed in^19, 20^). The direct target of mTORC1 on myogenesis has not been clearly established, but recent work has implicated mTORC1 in the regulation of MyoD protein stability, leading to a *miR*-1 dependent effect on myotube fusion^21^.

To determine the relevance of mTORC1 on muscle differentiation *in vivo* we have examined the effects of loss of TORC1 by both genetic and pharmacological approaches in the fruit fly, *Drosophila melanogaster*. In this study we present data supporting an essential developmental role of TORC1 in muscle development and/or integreity.

## Materials and Methods

### Tissue Culture and Myotube Formation

C2C12 cells were grown in High Glucose Dulbecco’s Modified Eagle’s Medium (DMEM; Sigma-Aldrich) supplemented with penicillin, streptomycin and glutamine (PSG; Life Technologies) and 10% Fetal Bovine Serum (Sigma-Aldrich). Once cells reached >90% confluence, differentiation media (2% Horse Serum from Sigma-Aldrich in DMEM with PSG) was added as previously described^22^ To determine when specific markers for differentiation were being expressed, cell lysates were prepared at time points between 0 and 15 days of differentiation. To determine the effects of rapamycin on differentiation, cells were treated every other day for 9 days with either vehicle alone (DMSO; Sigma-Aldrich), or 500nM rapamycin (Cayman chemicals) dissolved in DMSO. Cell lysates were prepared on day 9 of treatment, one days after the latest rapamycin administration. Cell lysates were generated by washing once with ice-cold PBS followed by the addition of 1 ml of QIAzol (Qiagen) and scraping into a 1.5ml microfuge tube. Lysates were stored at -80°C until RNA was purified.

### Quantitative Real Time PCR

RNA was extracted with the PureLink RNA mini kit (Life Technologies). 1 *µ*g of total RNA was used as a template to synthesize cDNA using the High Capacity Reverse Transcription Kit (Life Technologies). cDNA was added to *Power* SYBR Green PCR Master Mix in accordance with the manufacturer’s guidelines (Life Technologies) and qRT-PCR performed on a Roche Lightcycler. A series of control genes including *Gapdh, Rplp0, Actb* and *Rplp13a* were examined, and *Gapdh* was chosen as a control as it did not change across rapamycin concentrations or differentiation conditions. For a complete list of primers used, all purchased from IDT DNA, refer to Table 1. Relative expression was determined via the ΔΔCt method as previously described^23^.

**Table 1:**
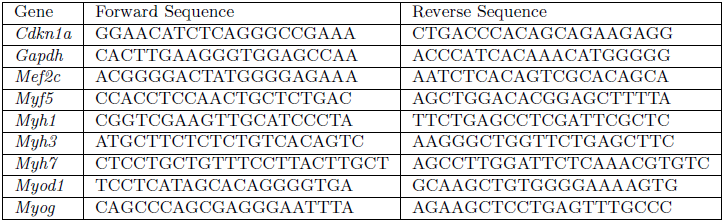
Forward and reverse primers used in qPCR experiments. All primers are based on mouse sequences.

### Protein Analysis

Cells were treated as indicated in the figure legend and then were lysed in RIPA buffer (50 mM Tris, pH 7.4, 0.25% sodium deoxycholate, 1% NP40, 150 mM sodium chloride, 1 mM EDTA, 100 uM sodium orthovanadate, 5 mM sodium fluoride and 10 mM sodium pyrophosphate) for 15 minutes on ice, then centrifuged for 15 minutes at 13 000 RPM at 4°C. Clarified lysates were loaded on SDS-PAGE gels, transferred and blotted using antibodies raised against MyoD (Pierce, cat # MA1-41017), pS6 (Serine 235/236, Cell Signaling cat # 2211), S6 (Cell Signaling cat # 2317), pAkt (Serine 473, Cell Signaling cat # 3787), Akt (Cell Signaling cat #2920). Antibody complexes were detected by anti-mouse and anti-rabbit fluorescent conjugated antibodies and visualized using an Odyssey image scanner and blots were quantified using the Odyssey software version 2.1 (LiCOR).

### Drosophila Stocks and Crosses

The stocks *w*^*1118*^, the three muscle GAL4 drivers (*24B*-GAL4, *Hand*-GAL4, *c179*-GAL4 and *Mef2*-GAL4), as well as both the *Raptor* and *Tsc1* UAS-shRNA TRiP lines used (See Table 2) were obtained from the Bloomington Stock Center (Bloomington, IN). All flies were raised at 25°C on standard corn meal food with the exception of the 18°C crosses for *24B*-GAL4. Rapamycin was added where indicated after fly food was cooled to below ~50° C. To prepare the crosses, virgin females were collected from each of the GAL4 driver strains. Ten virgin females were used per cross. Males with the appropriate genotype were chosen from each of the lines and crossed to male UAS-TRiP-shRNA lines for *Raptor* (3) or *Tsc1* (3) as well as a UAS-TRiP control which contains the genomic insertion site but no shRNA^24^. Flies were maintained in a humidified (50-60%) incubator at 25°C. A subset of experiments were also performed at 18°C. Ten days after each cross the F1 progeny began to eclose and adults were sorted according to phenotype and gender. During each sorting, the number of flies of each phenotype was recorded. The sorted flies were put into new vials, with males and females separated and with 5-10 flies in each vial. Progeny were stored at 25°C until at least 100 flies of each genotype had been collected. At least three independent replicates of each cross were performed.

**Table 2:** Fly stocks from Bloomington Stock Center used in this study.

| Name | Bloomington Stock |
| --- | --- |
| *Raptor* shRNA #1 | 31528 |
| *Raptor* shRNA #2 | 31529 |
| *Raptor* shRNA #3 | 34814 |
| *Raptor* shRNA #4 | 41912 |
| *Tsc1* shRNA #1 | 31039 |
| *Tsc1* shRNA #2 | 31314 |
| *Tsc1* shRNA #3 | 35144 |
| Control shRNA Line | 36304 |
| *Hand*-GAL4 | 48396 |
| *24B*-GAL4 | 1767 |
| *c179*-GAL4 | 6450 |
| *Mef2*-GAL4 | 27390 |
| *Mhc*-GAL4 | 38464 |

### Quantification of Dead Pupae

Twenty days after the *c179*-GAL4>UAS-shRNA-*Raptor* and *Mef2*-GAL4>UAS-shRNA-*Raptor* crosses were made any remaining adult or F1 progeny flies were emptied from the vials. The empty pupal cases were counted and the cases containing dead flies were counted. Pupal cases containing a dead fly were markedly darker in color than the empty cases and contained a visibly formed black, shrunken fly.

### Manual Assistance of Eclosure

In order to determine if flies were dying because they were too weak to eclose from their pupal cases or dead in their pupal cases for other reasons we manually removed the anterior puparial operculum under a dissecting microscope using fine forceps from stage ~12-13 pupae that were fully formed, but had not yet eclosed. This was accomplished by using a thin sheet of plastic on the inside of the vial on which the 3^rd^ instar larvae could form pupae. The sheet was then removed for imaging at various time points and to manually open the pupal cases then placed back into a fresh vial for incubation at 25°C. Using this method allowed for the rescue of 3 *Mef2*-GAL4>UAS-*Raptor*-shRNA adults that were too weak to begin to eclose, but with assistance of the removal of the operculum could get out of the case, inflate their wings and appeared morphologically normal.

### Climbing Assay

To perform the climbing assay flies were tapped to the bottom of a vial and a stopwatch was started simultaneously. The stopwatch was stopped each time a single fly from the group in the vial climbed to a mark at 4cm on the side of the vial. A separate time was recorded for each fly in the vial. This assay was first performed within 3 days post eclosure and repeated every ~30 days for a total of 3 trials.

### Statistics

Statistical analyses were performed using the R statistical package (version 3.1.0)^25^. Prior to performing ANOVA analyses, normality was assessed by Shapiro-Wilk tests and equal variance was tested using Levene’s tests (from the car package (version 2.0-20)^26^). If both these assumptions were met (p>0.05) an ANOVA was performed. If either of these assumptions were not met, a Kruskal-Wallis test was performed. If either of those omnibus tests reached significance, then Student’s *t*-tests or Wilcoxon Rank Sum Tests were performed as indicated, followed by an adjustment for multiple comparisons using the method of Benjamini and Hochberg^27^. Statistical significance for the manuscript was set at a *p* or *q*-value of < 0.05. The investigators were blinded to the genotype of the crosses until analysis. For barplots, data represents the mean +/- the standard error of the mean. All raw data, analyzed data and code used to analyze the data and generate figures is available at http://bridgeslab.github.io/DrosophilaMuscleFunction/^28^

## Results and Discussion

### Rapamycin Inhibits Differentiation of Muscle Cells in Culture

To determine the order in which myogenic markers are induced during myogenesis, we performed a time course experiment in C2C12 cells. We generated cell lysates at various time points between 0 and 15 days of the differentiation process and performed qRT-PCR to measure transcripts of known differentiation markers including *Myf5*, *Myog*, *Mef2c*, *Cdkn1a*, and *Myod1* as well as the major myosin heavy chain genes in these cells *Myh1, Myh3* and *Myh7*. We observed that transcripts for *Myf5*, *Myog*, *Cdkn1a*, and *Myod1* are increased early in the process (~ day 2) and continue to increase throughout development with large increases in *Mef2c* not occurring until approximately day 5 and *Myh1* increasing around day 7 (Figure 1a). This is consistent with previous observations of the transcriptional changes associated with muscle differentiation of cells in culture^29, 30^.

**Figure 1.**
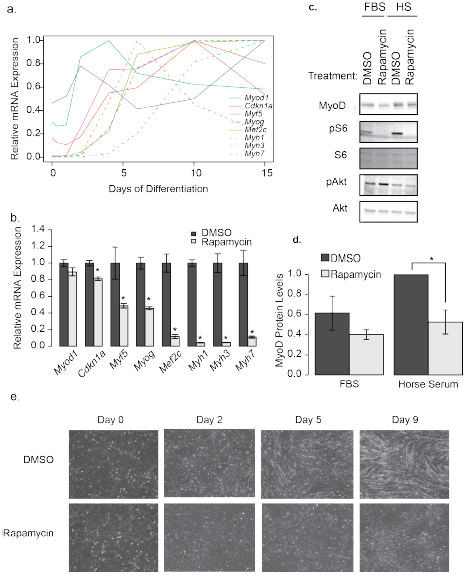
Rapamycin blocks C2C12 differentiation. a) The order of appearance of myotube differentiation markers over the course of 15 days in differentiation media only. This is representative of three independent experiments. b) Differences in differentation marker transcripts when treated with DMSO (vehicle) or 500nM rapamycin for 9 days, throughout the differentiation protocol. The most recent rapamycin administration was 1 day prior to cell lysis. These data represent the average of three wells, from a representative experiment (n=3). Transcripts from both a) and b) were measured by qRT-PCR and normalized to *Gapdh*. c) Representative western blot analysis of C2C12 cells treated for 4h with Differentiation Media (2% Horse Serum; HS) or left in growth media (10% FBS) in the presence of 500 nM Rapamycin or DMSO. d) Quantification of the blots in c (n=4 independent experiments). Protein phosphorylation is presented as the intensity of the phosphospecific antibody band relative to that proteins total protein band. e) Images of morphological changes in C2C12 myoblasts in response to 10 days of DMSO or rapamycin treatment througout the differentiation protocol (500nM). Asterisks indicate p<0.05. Data represents mean +/− standard error of the mean.

Next we wanted to determine if rapamycin, a drug known to inhibit TORC1 signaling, had any effects on gene expression during differentiation (Figure 1b and c). Treatment with rapamycin throughout the differentiation protocol caused significant reductions in mRNA transcript levels detected for all differentiation markers measured (p<0.05), with the exception of *Myod1* (Figure 1b) and prevented the formation of myotubes (Figure 1c). We did not observe any fused myocytes in the rapamycin treated cells, suggesting that the earliest rapamycin-sensitive event is prior to myocyte fusion, which results in impaired myotubule formation. This is consistent with previous studies examining the effects of rapamycin on myoblast differentiation^9, 11, 31–33^.

Since *Myodm*RNA levels were unchanged, we next tested whether MyoD protein levels are altered by rapamycin treatment. We added the differentiation media for 4h in the presence of DMSO or rapamycin and observed that rapamycin reduced MyoD protein levels in by 47% (Figures 1c-d). These data are consistent with the hypothesis that one role of mTORC1 in differentiation is through the stabilization of MyoD as previously suggested^21^, though whether there are other mTORC1 targets in early differentiation is not clear. Since the primary effect of miRNA-1 is on myotube fusion, it is likely that there are other mTORC1 dependent effects, as the morphological changes prior to myotube fusion are also disrupted by rapamycin^11^.

We also observed elevations in mTORC1 activity 4h after the transition to differentiation media, as shown by increased S6 (Figure 1c). These data suggest that activation of mTORC1 signaling occurs during differentiation, consistent with previous reports^11^. Furthermore, this activation is independent of Akt signaling, which was actually decreased during the transition from 10% FBS to 2% horse serum. Morphologically, there was a complete lack of fused myotubes in the rapamycin treated cells at all time points observed (Figure 1e).

### Muscle Specific Knockdown of *Raptor* Leads to Late Pupal Lethality in Drosophila

In order to study the role of TORC1 signaling on muscle development *in vivo*, we manipulated dTORC1 function in the model organism *Drosophila melanogaster* (fruit flies). First, we tested whether inhibition of the dTORC1 pathway affected the development of these flies. As previously reported, high doses of rapamycin prevents egg laying in females^16^. We performed dose curves and found that at much lower doses (EC50 of ~860 nM), although eggs could be seen in the vials, there was a complete absence of pupae and adult flies (Supplementary Figure 1). At these lower doses, there was no obvious distinction between inhibition of pupal lethality and prevention of fly eclosure, ie there was no observable dose in which pupae survived but flies were unable eclose. These data suggest that rapamycin inhibits fly muscle development, similar to what has been observed in mice^12^. It also supports studies showing that whole animal knockout of *Raptor* leads to developmental lethality in several model organisms13, ^15, 34^.

To look specifically at the role of dTORC1 in muscle, we knocked down either *Tsc1* or *Raptor* to generate constitutive gain and loss of function of mTORC1 activity in fly muscles using the GAL4-UAS system^35^. We used several GAL4drivers that drove expression of the UAS shRNA cassettes in both skeletal muscle and cardiac muscle. We targeted skeletal muscle using *24B*-GAL4, *C179*-GAL4, and *Mef2*-GAL4 drivers, while cardiac muscle was targeted using the heart specific *Hand*-GAL4 driver. To minimize potential off target effects, three different shRNAs were used from the Harvard shRNA TRiP collection for each of the two genes (*Raptor* and *Tsc1*).

First, we crossed heterozygous, balanced *24B*-GAL4/TM3, *Sb* flies with heterozygous, balanced UAS-shRNA/TM6B transgenic flies. The flies inheriting both balancer chromosomes had decreased viability and were excluded from the analysis. A control strain, expressing no shRNA had a modest decrease in the number of flies with the TM3,Sb/Control genotype (47% of flies of this genotype, with an expected ratio of 50%, n=537 flies, p=1 by Fisher’s test). Progeny from crosses using the *Hand*-GAL4 driver appeared in roughly equal ratios (Figure 2a), indicating there is no obvious effect of manipulating dTORC1 in cardiac cells. Similarly, *24B*-GAL4 driven expression of *Tsc1* shRNA had no significant effect on viability. However, when the *24B*-GAL4 driver was used to express *Raptor* shRNA, there was a dramatic decrease in the number of eclosed flies (Figure 2b). This indicates that *24B*-GAL4 driven expression of *Raptor* shRNA is lethal at some point prior to eclosure. Similarly, another muscle specific driver, *c179*-GAL4 crossed to heterozygous UAS-*Raptor*-shRNA/TM6B resulted in a reduced number muscle-specific *Raptor* knockdown flies (i.e. *c179*-GAL4>UAS-*Raptor*-shRNA), although in this case some flies expressing UAS-*Raptor*-shRNA were able to eclose (Figure 2c).

**Figure 2.**
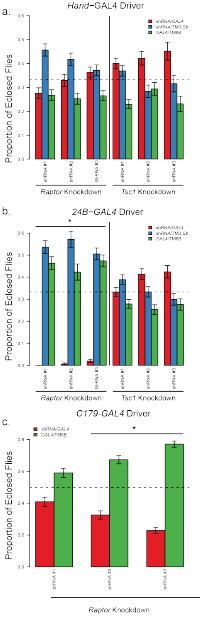
Skeletal muscle specific Raptor knockdown causes lethality. a) Proportion of progeny born from a *Hand*-GAL4/TM3, Sb x shRNAi/TM6B, *Tb, Hu*. The progeny that are TM6B/TM3 were excluded due to known reduced viability of flies with balancer chromosomes so the expected ratios (as indicated by the dotted line) are 0.33. Knockdown flies are shown in red throughout. b) Proportion of progeny born from a *24B*-GAL4/TM3, *Sb* x shRNA/TM6B, *Tb, Hu* cross. c) Proportion of progeny born from a *c179*-GAL4/*c179*-GAL4 x shRNAi/TM6B, *Tb, Hu* cross. In this case half the progeny should be knockdown, so the expected ratio is 0.5. Asterisks indicates p<0.05, testing the hypothesis that the knockout flies eclose at less than the expected proportions. Error bars indicate sampling standard error, with >195 flies examined for each cross.

We next attempted to rescue the lethality in *24B*-GAL4>UAS-*Raptor*-shRNA flies by lowering the temperature of the cross to 18 °C. Colder temperatures decrease GAL4 expression in driver lines^35^. Decreasing the temperature to 18 °C did not rescue the lethality of the *24B*-GAL4/UAS-*Raptor*-shRNA flies, and the birth rates of the two control genotypes were congruent with birth rates at 25° C (Supplementary Figure 2).

To test for the stage at which these flies fail to eclose, we next used *c179*-GAL4 and *Mef2*-GAL4, which drives expression late in muscle development^36^, and repeated the studies at 25 °C. As a control, we used a fly line that was identical to the TRiP fly lines, but did not have a shRNA inserted (see Table 2). All flies where *Raptor*-shRNA was driven by *Mef2*-GAL4 died prior to eclosion (see Figure 3a). There was partial lethality in the three the c179-GAL4 mediated *Raptor* knockdown flies (q-value < 0.005 for those shRNA strains, with a 73-92% decrease in the number of flies depending on the strain, see Figure 3b). These results indicate that the c179-GAL4 driver is less efficient at mediating *Raptor*-specific lethality than the *Mef2*-GAL4 and the *24B*-GAL4 drivers.

**Figure 3:**
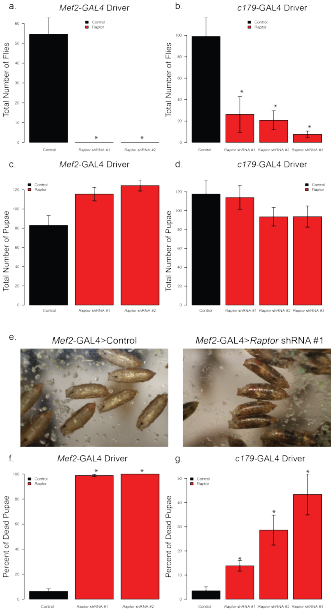
Muscle-specific Raptor knockdown flies die around pupal stage ~12-13 but pre-eclosure. The graph depicts the birthrates of the progeny from crosses of homozygous GAL4driver lines with homozygous *Raptor* shRNA transgenic flies or a control TRiP line. Panels a, c and f indicate *Mef2*-GAL4 driven knockdowns while b, d and g indicate *c179*-GAL4mediated knockdowns. a) and b) show the total number of flies eclosed; c) and d) indicate the total number of pupae after 20 days, and f) and g) show the percentage of dead pupae. Panel e) shows a representative example of dead flies, still within their pupal cases for *Mef2*-GAL4>UAS-*Raptor*-shRNA #1. Asterisks indicate p<0.05 by ANOVA followed by Dunnett’s test (b, c and d) or Kruskal-Wallis tests then Wilcoxon rank-sum tests followed by an adjustment for multiple comparisons (a, f and g). Each of these analyses describe the average 5-9 independent crosses, with error bars indicating standard error of the mean between replicate crosses.

### Muscle *Raptor* Knockdown Flies Fail to Eclose from Pupae

To determine at which point prior to eclosure the *Raptor* knockdown flies die, we examined the pupal cases on the sides of the vials from the *cl79*-GAL4>UAS-*Raptor*-shRNAcrosses and the *Mef2*-GAL4>UAS-*Raptor*-shRNAcrosses. Twenty days after the crosses were prepared both the empty pupal cases and the cases containing dead flies were counted. We observed no significant differences in the total number of pupal cases from either of these crosses (Figure 3c-d, p=0.416 and p=0.066 from ANOVA respectively). In fact, we observed a slightly increased number of pupae from the *Mef2*-GAL4 > UAS-*Raptor*-shRNA crosses. These data support the hypothesis that lethality occurs after pupal development.

We next visually examined the pupal cases for the presence dead flies (Figure 3e). After blind scoring, we noted that for the *Mef2*-GAL4 driven *Raptor* knockdown nearly 100% of the pupal cases contained dead flies using two different anti-*Raptor* shRNA lines (15 fold more dead pupae than controls; Figure 3f). There was also a significant number of dead flies in pupal cases from the *cl79*-GAL4*>*UAS*-Raptor*-shRNAcrosses (Figure 3g). Although the absolute number of dead pupae was variable among the shRNA-*Raptor* lines using c179-GAL4, in all cases the percentage of dead flies in pupal cases was significantly greater than controls (Figure 3g). These results demonstrate that *Raptor* knockdown in skeletal muscle produces lethality after pupal development, but prior to eclosure.

### Lethality of *Raptor* knockdown in skeletal muscle is due to an inability to eclose from the pupal case

To test whether the muscle *Raptor* knockdown-mediated lethality is due to a muscle weakness that prevents eclosure, we first carefully examined fly morphology within pupal cases. As shown in Figure 4a ten days after the cross a fully formed fly is visible within the pupal case and looks morphologically similar to control flies. By day 14 the control flies have completely eclosed leaving only empty pupal cases while the *Mef2*-GAL4>*Raptor* flies are still in the pupal case. By day 20 the flies have started to decompose and appear dark as in Figure 3e.

**Figure 4.**
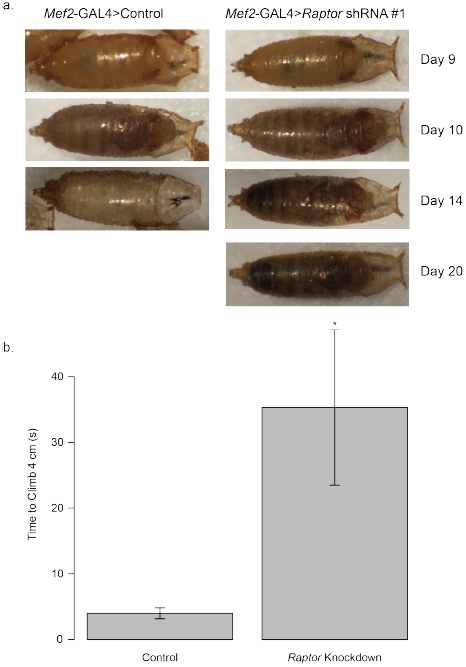
Mef2-GAL4Driven *Raptor* Flies Cannot Eclose from Pupal Cases. a) Flies with *Mef2*-GAL4 driven knockdown of the control shRNA or *Raptor* shRNA #1 were examined from pupal stage 10 till after eclosure. By day 14 after the cross, control flies had completely eclosed but *Raptor* knockdown flies remained inside the pupal cases. b) On day 10, (approximately pupal stage 12) pupal cases were cut open at the operculum to assist eclosure, then after 4 days, climbing assays were performed on escaping flies. The *Mef2*-GAL4 *> Raptor* shRNA flies that eclosed with assistance had significantly impaired climbing. Asterisk indicates p<0.005 by Wilcoxon Rank Sum test. Data represents mean +/− standard error of the mean.

To determine if this is due to an inability of the fly to exit the pupal case, we gently opened 5 pupal cases by removal of the operculum at day 10 (approximately stage 12-13 pupae) from *Mef2*-GAL4>*Raptor* knockdown flies to assist in eclosure. In 4 out of 5 cases the flies eclosed successfully with 3 of these animals surviving >3 weeks. To validate that these eclosure-assisted flies had muscle weaknesses we performed climbing assays as shown in Figure 4b. The *Mef2*-GAL4>*Raptor* flies exhibited dramatically reduced climbing ability as compared to controls indicating muscle weakness (p=0.0025 by Wilcoxon Rank Sum Test).

### Effects of Muscle-Specific *Raptor* Knockdown on Longevity

We next turned our attention to the few flies that survived from the *c179*-GAL4 cross. The lifespan of these *Raptor* knockdown flies was measured to determine the effects of dTORC1 suppression on longevity. When *Raptor* was knocked down in skeletal muscle using the *c179*-GAL4 driver, a large proportion of the flies that successfully eclosed died shortly afterwards. Interestingly, among the flies that survived, they generally had normal lifespan (see Figure 5).

**Figure 5.**
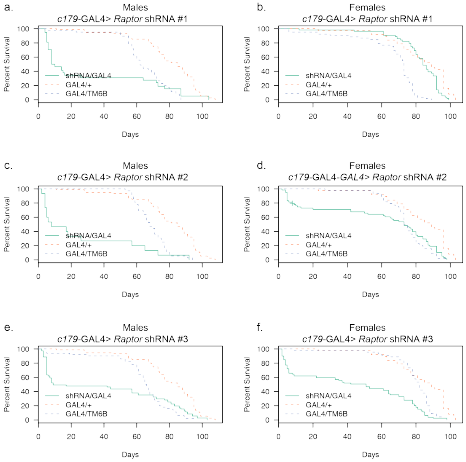
Lifespan of *C179*-GAL4 Driven *Raptor* Knockdown Flies. Dashed lines indicate two control strains. Each panel shows a control of *C179*-GAL4 crossed to the control shRNA strain, as well as the balancer containing progeny of the *C179*-GAL4 homozygotes crossed to the heterozygous UAS-*Raptor* shRNA/TM6b flies. *C179*-GAL4>*Raptor* shRNA #3 had the strongest effect on lifespan for both males and females, but all three *Raptor* shRNA constructs significantly decreased survival.

### Effects of Muscle Specific *Raptor* Knockdown on Muscle Function

To study the effects of dTORC1 suppression on muscle function, a climbing assay was performed on the *Raptor* knockdown flies driven by the *c179*-GAL4 driver at several ages. Progeny from each cross were individually timed for how long it took them to climb 4 cm up the side of the vial. The average times for each cross are shown in Figure 6. The results indicate that dTORC1 suppression leads to reduced muscle function in the flies that eclose even very early, consistent a developmental problem in myogenesis. Notably, these problems persist throughout the lifespan of the fly, even in those animals that reach adulthood and have an average lifespan. Also interesting, is that there was a correspondence between the efficiency of the shRNA strain to cause lethality and its effects on climbing ability, indicating a potential gene-dosage effect on both of these phenotypes. This is consistent with other work in flies showing a correlation between climbing and lifespan^37–40^, indicating that muscle strength and aging are often linked, as is observed in humans^41–43^.

**Figure 6.**
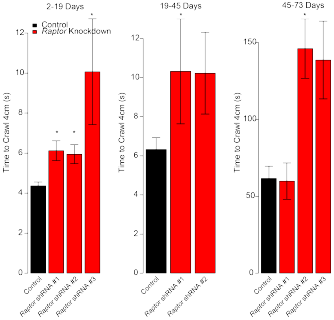
*c179*-GAL4 *Driven Raptor* knockdown flies have reduced climbing rate. Average climbing rate as measured during three age range intervals (in days) for *c179*-GAL4 driven *Raptor* knockdown flies. Asterisk indicates p<0.05 based on a Wilcoxon Rank-Sum test relative to the control flies, and adjusted for multiple observations. Note that the different abscissa indicates age-related slowing of climbing speed. Data represents mean +/− standard error of the mean.

## *Mhc*-GAL4 Driven Raptor Knockdown Does Not Result in Early Muscle Defects

In order to evaluate the effects of *Raptor* knockdown later during differentiation, we next utilized an *Mhc*-GAL4 driver. *Mhc* expression occurs quite late in the differentiation process relative to *Mef2* in differentiating C2C12 cells (Figure 1a). In contrast to the other, earlier GAL4 lines, we did not observe any defects in eclosure with *Raptor* knockdown using the *Mhc*-GAL4 driver (Figure 7a). We then evaluated the eclosed flies for climbing activity, and did not observe any significant differences between these flies and control flies, although there was a slight trend towards decreased climbing activity (Figure 7b). We then evaluated the lifespan of these flies and found that in spite of no significant changes in observed birth rates, or climbing ability, both male and female flies tended to die earlier than control flies (Figure 7c). These results indicate that raptor continues to play a role in muscle function after development and eclosure.

**Figure 7:**
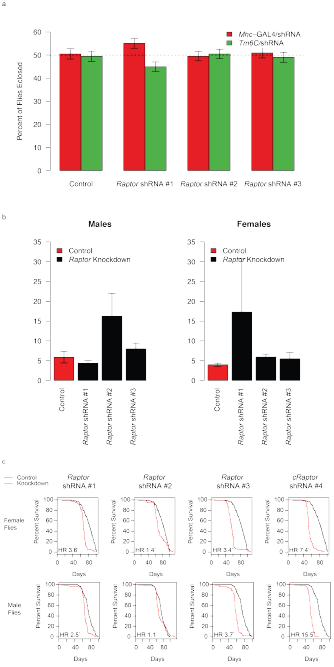
*Mhc*-GAL4 *Raptor* knockdown flies have no detectable decreases in viability or climbing ability. a) Birth rates for crosses of *Mhc*-GAL4/*TM6c* vs homozygous shRNA strains. The expected birth ratio of this cross is equal numbers of progeny and is indicated by the dashed line. b) Climbing activity for flies described in a, measured between 2 and 19 days of age. c) Longevity analysis for *Mhc*-GAL4 > *Raptor* knockdown flies. Four shRNA strains are shown compared to the control shRNA strain crossed to *Mhc*-GAL4 (n>98 for each gender/cross). HR indicates the hazard ratio, asterisk indicates q<0.05. Data represents mean +/− standard error of the mean.

## Discussion

Several previous reports have implicated mTORC1 activity or activation as a necessary step in myogenesis *in vitro*^9–11,21,33^. Our data is consistent with these findings. We provide data in support of the hypothesis that destabilization of MyoD with rapamycin treatment is an early inciting event in the inhibition of myogenesis, occurring within hours of treatment, consistent with previous observations^21^. Although mTORC1 is the primary target of rapamycin, studies using rapamycin-insensitive analogs of mTOR have suggested there may be alternative rapamycin-sensitive targets of rapamycin at other stages of myogenesis^11,20,33^.

At the end of the differentiation process, we did not observe any changes in *Myod* mRNA levels, but we did observe decreases in several other early differentiation targets, including *Myog*, *Myf5*, and *Cdkn1a* are all blocked by rapamycin. Since *Mef2c* is downstream of *Myog*, reductions in *Mef2c* levels are likely due to defects upstream of *Myog*^44^. The decreases in the mRNA levels at the end of the study for *Myog*, *Myf5*, *Mef2c* and *Cdkn1a* are likely reflective of undifferentiated cells, and may not be direct mTORC1 targets. Although these data do not preclude the possibility of other unknown factors, our observations support the hypothesis that mTORC1 is required for MyoD stability, which is then required for activation of the remainder of the myogenic program.

To extend these *in vitro* findings into an *in vivo* system we have examined a panel of muscle-specific GAL4 drivers to knock down the *Raptor* gene in flies. We observed a complete or near-complete inability of flies to eclose with *Mef2 and 24B* drivers along with partial lethality using the *c179* driver, but importantly, no lethality with the *Mhc* driver. All three of *24B*-GAL4^45^, *Mef2*-GAL4^46^ and *c179*-GAL4^47,48^ are reported to be expressed in wing disks as well as muscle and *Mhc* has been shown to be expressed in the developing embryo in addition to differentiated muscle^49^. A complete evaluation of the precise timing of activation of these drivers was not performed in this study, but one possibility is that *Raptor* is required for efficient muscle development at a stage corresponding to the *Mef2/24B* promoter activation, but is no longer required by the time *Mhc* is expressed. This hypothesis is supported by the finding that in C2C12 cells, *Mhc* genes are elevated after *Mef2c* and the other myogenic transcription factors (Figure 1a). Furthermore, mRNA profiling studies of wing disc derived cells lines show expression of *Mef2* but not *Mhc* in these developing organs^50^. Alternately, it is possible that the differences observed between muscle drivers are due to differences in knockdown efficiency or different anatomical locations in which these drivers are active. A previous way to reduce dTORC1 signaling is to overexpress *Tsc2* a negative regulator of dTORC1 signaling. Using the *24b*- GAL4 transgene to drive UAS-*Tsc2* expression, Kapahi and colleagues showed that these flies had reduced lifespan, consistent with our findings, although they did not report any eclosure defects^51^.

As shown in Figure 4, we are able to rescue the pupal lethality of these flies by assisting with their eclosure from pupal cases, but even when the flies emerge, they are noticeably weaker. This suggests that there may be a developmental defect or muscle maintenance defect *Mef2*-GAL4 > *Raptor* knockdown flies, and the observed lethality is most likely due to an inability to emerge from pupal cases due to weakened muscle strength.

These findings are somewhat consistent with mouse studies in which human skeletal actin-driven (*ACTA1*) Cre expression drove the knockout of muscle *Rptor*. These mice were observed to be weaker than littermate controls, and prone to early death^52^, similar to our observations of the *c179*-GAL4 and *Mhc*-GAL4 driven *Raptor* knockout flies. In the mouse model *Rptor* is not expected to be knocked down until post-differentiation, as *Acta1* is expressed late in myogenesis, and not at all in satellite cells^53^, so it is probable that these mice die of an alternative muscle-specific defect later in life, and not a developmental myogenic defect. In *c179*-GAL4 driven *Raptor* knockdown flies we observed a critical period of about 20 days after eclosure during which the *Raptor* knockdown flies are still prone to early death. Furthermore, even outside of the context of reduced viability/climbing ability *Mhc*-GAL4 driven *Raptor* knockdown flies still died earlier than control flies (Figure 7c). The *ACTA1-Cre* driven *Rptor* knockout studies did not evaluate mTORC1-dependent myogenesis in mice. Another study implicated mTORC1 in the differentiation of ES cells into satellite cells, a process which is likely upstream of our model system^54^. Together these results implicate mTORC1 as essential at multiple steps of myogenesis and maintenance of muscle function in both flies and mice.

## Acknowledgements

The authors would like to thank the members of the Bridges, Bissler and Reiter labs for insightful discussions and C. Valdez for assistance with animal husbandry. We would like to thank Markus Ruegg (University of Basel) for helpful discussions regarding this work. The qPCR instrumentation used in this study was provided by the Molecular Resource Center at UTHSC, and we would like to thank William Taylor and Felicia Waller for their assistance. This study used stocks obtained from the Bloomington Stock Center (funded by NIH Grant P40OD018537). This work was supported in part by a Dean’s Neurology Support Fund to LTR and the UTHSC Drosophila Transgenic Core.

